# Protein-bound molybdenum cofactor is bioavailable and rescues molybdenum cofactor-deficient *C. elegans*

**DOI:** 10.1101/2020.10.08.332338

**Authors:** Kurt Warnhoff, Thomas W. Hercher, Ralf R. Mendel, Gary Ruvkun

## Abstract

The molybdenum cofactor (Moco) is a 520 dalton prosthetic group that is synthesized in all domains of life. In animals, four oxidases (among them sulfite oxidase) use Moco as a prosthetic group. Moco is essential in animals; humans with mutations in genes that encode Moco-biosynthetic enzymes display lethal neurological and developmental defects. Moco supplementation seems a logical therapy, however the instability of Moco has precluded biochemical and cell biological studies of Moco transport and bioavailability. The nematode *Caenorhabditis elegans* can take up Moco from its bacterial diet and transport it to cells and tissues that express Moco-requiring enzymes, suggesting a system for Moco uptake and distribution. Here we show that protein-bound Moco is the stable, bioavailable species of Moco taken up by *C. elegans* from its diet and is an effective dietary supplement, rescuing a *C*. *elegans* model of Moco deficiency. We demonstrate that diverse Moco:protein complexes are stable and bioavailable, suggesting a new strategy for the production and delivery of therapeutically active Moco to treat human Moco deficiency.

## Introduction

The molybdenum cofactor (Moco) is an ancient coenzyme that was present in the last universal common ancestor and that continues to be synthesized in all domains of life (Weiss et al. 2016; Zhang and Gladyshev 2008). Moco is a pterin-based organic prosthetic group that is comprised of a C6-substituted pyrano ring, a terminal phosphate, and a dithiolate group binding to molybdenum (**Fig. 1A**) (Rajagopalan and Johnson 1992). In humans and other animals, Moco is required for the activity of 4 enzymes: sulfite oxidase, xanthine oxidase, aldehyde oxidase, and mitochondrial amidoxime reducing component (Schwarz et al. 2009). There are 2 forms of eukaryotic Moco, the sulfite oxidase form and the xanthine oxidase form (**Fig. 1A)**. These Moco species differ in the third Mo-S ligand which is provided either by an enzyme-derived cysteine residue (sulfite oxidase form) or an inorganic sulfur (xanthine oxidase form) (Schwarz et al. 2009). The xanthine oxidase form of Moco is synthesized from the sulfite oxidase form via the enzyme Moco sulfurase (**Fig. 1A**) (Bittner et al. 2001).

**Figure 1:**
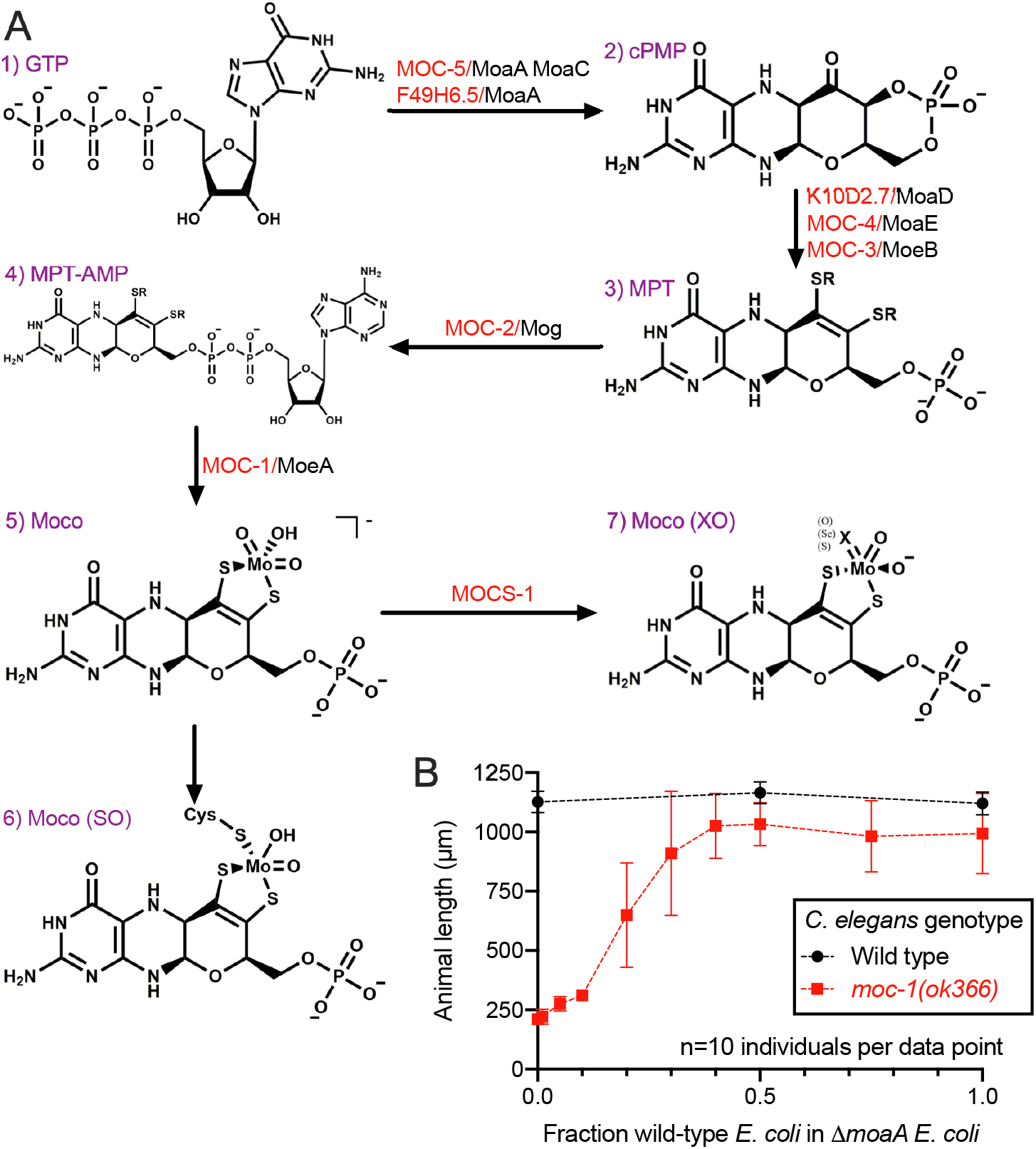
*C. elegans* acquires Moco from dietary *E. coli.* (A) *C. elegans* Moco biosynthesis pathway (red) and orthologous enzymes in *E. coli* (black) are displayed. Moco and its biosynthetic intermediates are displayed (purple): GTP is guanosine triphosphate (1), cPMP is cyclic pyranopterin monophosphate (2), MPT is molybdopterin (3), MPT-AMP is MPT-adenine monophosphate (4), Moco is the molybdenum cofactor (5), Moco (SO) is the sulfite oxidase form of the molybdenum cofactor (6), and Moco (XO) is the xanthine oxidase form of the molybdenum cofactor (7). *C. elegans* Moco sulfurase (MOCS-1) has no clear homolog in *E. coli,* although *xdhC* is the likely functional analog (Neumann et al. 2007). (B) Wild-type and *moc-1(ok366) C. elegans* were synchronized at the L1 stage and cultured on mixtures of wild-type *E. coli* (synthesizes Moco) and Δ*moaA E. coli* (cannot synthesize Moco) for 72 hours. The Y axis shows animal length (µm), where 1,000µm roughly corresponds with fertile adulthood and 250µm roughly corresponds to the L1 stage. Average and standard deviation are displayed for each condition analyzed. Sample size (n) was 10 individual animals assayed for each condition.

Both forms of Moco are synthesized by a highly conserved biosynthetic pathway (**Fig. 1A**) (Mendel 2013). The genes necessary for Moco biosynthesis were first elucidated by genetic studies of chlorate resistance in bacteria (MacGregor 1975). The importance of Moco biosynthesis to human health is highlighted by Moco deficiency (MoCD), a rare inborn error of metabolism. MoCD is caused by loss-of-function mutations in genes encoding any of the human Moco-biosynthetic enzymes and results in severe neurological dysfunction and neonatal lethality (Reiss and Hahnewald 2010; Huijmans et al. 2017). MoCD patients with mutations in MOCS1 (orthologous to bacterial *moaA* and *moaC*) can be treated with cyclic pyranopterin monophosphate (cPMP), a stable intermediate in Moco biosynthesis immediately downstream of MOCS1 (Veldman et al. 2010). However, cPMP treatment is not effective for patients with mutations in any of the downstream Moco-biosynthetic enzymes. Purification and delivery of mature Moco would be an ideal therapeutic strategy for treating all forms of MoCD, however free Moco is too unstable and oxygen-sensitive to be purified and therapeutically administered (Johnson et al. 1980; Mendel 1983). Furthermore, it is unclear whether mature Moco can cross cellular membranes.

Genetic evidence demonstrates that the nematode *C. elegans* retrieves Moco as well as cPMP from its bacterial diet (Warnhoff and Ruvkun 2019). However, nothing was known about the biochemical mechanism of Moco transfer between these 2 highly divergent organisms. Here we propose that Moco bound to protein is the stable and bioavailable Moco species that is harvested by *C. elegans.* We demonstrate that supplementation of purified protein-bound Moco rescues the lethality of Moco-deficient *C. elegans* feeding on Moco-deficient *E. coli*. We show that Moco bound to diverse Moco-containing proteins originating from bacteria, algae, fungi, and mammals, is bioavailable to *C. elegans,* and that this supplementation does not require Moco biosynthetic enzymes in *C. elegans* or its bacterial diet. This work suggests future mammalian therapeutic studies of supplemental protein-bound Moco and highlights the existence of a pathway for Moco transport.

## Results and Discussion

### *C. elegans* acquires Moco from dietary *E. coli*

Due to its instability, Moco has long been thought to be synthesized and utilized cell autonomously. So far only *C. elegans* has been described to have 2 pathways by which it can obtain Moco: endogenous Moco biosynthesis from GTP or dietary uptake of Moco (Warnhoff and Ruvkun 2019). Moco-biosynthetic enzymes are conserved in all domains of life; in *C. elegans* these enzymes are encoded by the *moc* genes that mediate sequential steps in Moco biosynthesis **(Fig. 1A)**. Using mutations in the *C. elegans moc* genes (i.e. the *moc-1(ok366)* null mutation) Moco biosynthesis can be interrupted in all cells. In the laboratory, *C. elegans* feed on a monoculture of *E. coli*. Thus, we can also use mutations in any of the genes of the *E. coli* Moco-biosynthetic pathway to eliminate dietary Moco (i.e. theΔ*moaA* null mutation). Either endogenous Moco synthesis in *C. elegans* or Moco produced by the diet *E. coli* and then consumed by *C. elegans* can support growth, development, and reproduction of *C. elegans*. However, when *C. elegans* cannot synthesize their own Moco and cannot obtain Moco from their diet, they arrest larval development and die due to inactivity of sulfite oxidase, the key Moco-utilizing enzyme in animals (Warnhoff and Ruvkun 2019).

To test how much wild-type, Moco-producing bacteria was required to support growth and development of *C. elegans* defective in Moco biosynthesis, we mixed wild-type and Δ*moaA* mutant *E. coli* at various ratios and tested for the ability of these mixtures to support the viability of *moc-1* mutant *C. elegans.* We found that a substantial fraction (about 30%) of the *E. coli* diet needed to be wild type (Moco producing) to support growth and development of *moc-1* mutant animals (**Fig. 1B)**.

### Moco bound to diverse proteins is taken up and used by *C. elegans*

We hypothesized that *C. elegans* harvest bacterial Moco that is bound within the *E. coli* Moco-utilizing enzymes. *E. coli* YiiM (*Ec*YiiM) is one such Moco-utilizing enzyme and mediates the reduction of N-hydroxylated substrates (Namgung et al. 2018; Kozmin et al. 2008). To test whether Moco bound to *Ec*YiiM can be absorbed by *C. elegans*, we purified recombinant *Ec*YiiM protein from *E. coli* and used it to supplement the diet of Moco-biosynthetic mutant *C. elegans* feeding on Moco-deficient *E. coli,* growth conditions that would otherwise result in 100% larval arrest and death. Consistent with the model that *C. elegans* harvests Moco from *E. coli* Moco-utilizing enzymes, *moc-1* mutant animals grown on Moco-deficient *E. coli* grew and developed well when their diet was supplemented with *Ec*YiiM-bound Moco. (**Fig. 2A,B**). Thus, *Ec*YiiM-bound Moco is bioavailable and can support the viability of otherwise Moco-deficient *C. elegans*.

**Figure 2:**
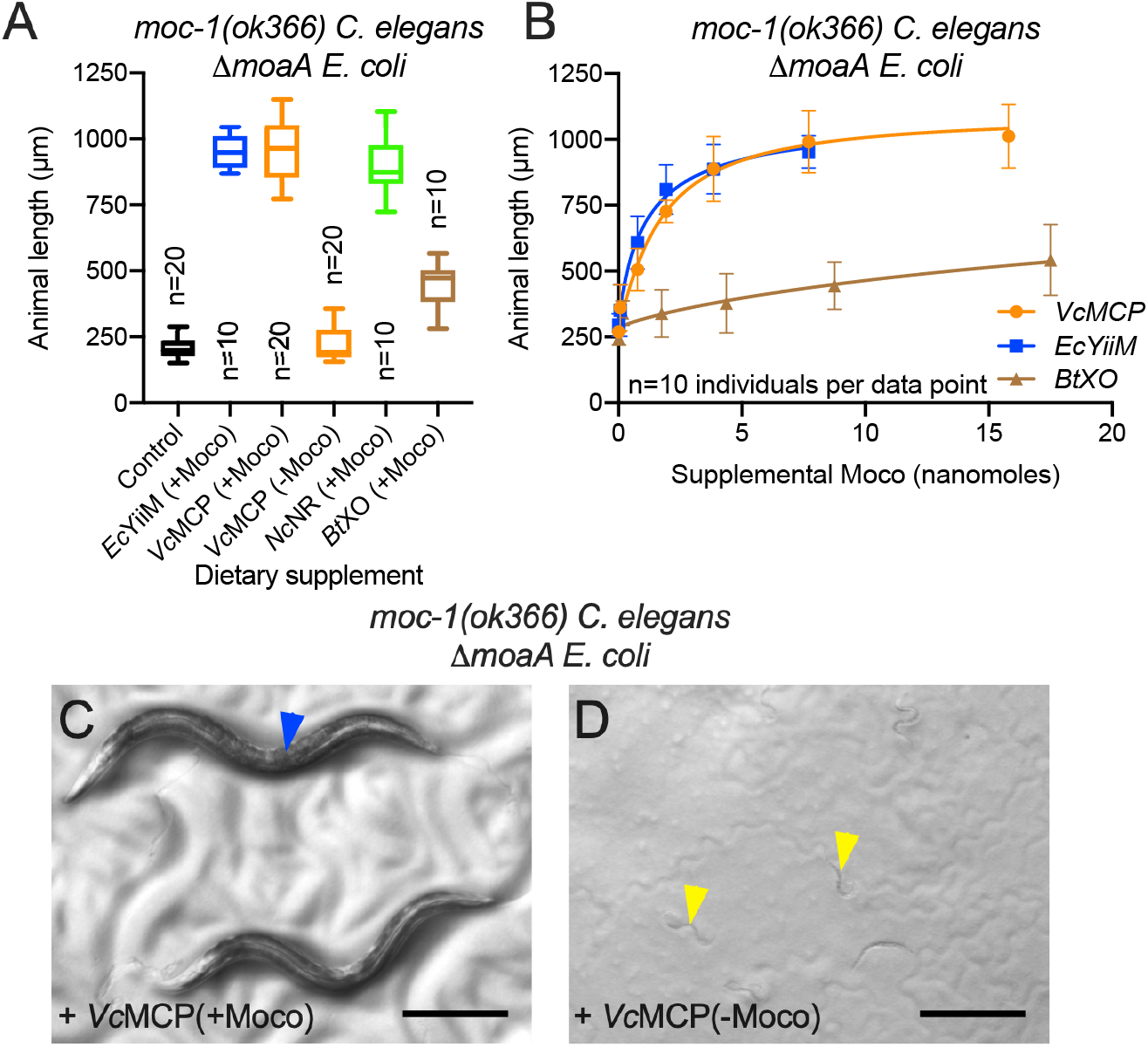
*C. elegans* uses Moco from diverse Moco-containing proteins. (A) *moc-1(ok366)* mutant *C. elegans* were synchronized at the L1 stage and cultured for 72 hours on Δ*moaA E. coli* supplemented with Moco bound to *Escherichia coli* YiiM (*Ec*YiiM), *Volvox carteri* Moco carrier protein (*Vc*MCP), *Neurospora crassa* nitrate reductase (*Nc*NR), or bovine xanthine oxidase (*Bt*XO), or equivalent amounts of *Vc*MCP purified from bacteria that cannot synthesize Moco (-Moco). *Ec*YiiM, *Vc*MCP, and *Nc*NR (+Moco) each contained 7.7 nanomoles of Moco while *Bt*XO (+Moco) contained 8.8 nanomoles of Moco. Box plots display the median, upper, and lower quartiles while whiskers indicate minimum and maximum data points. Sample size (n) is displayed for each experiment. (B) *moc-1(ok366)* mutant *C. elegans* were synchronized at the L1 stage and cultured on Δ*moaA E. coli* supplemented with variable amounts of Moco bound to *Ec*YiiM (0.0077, 0.077, 0.77, 1.93, 3.85, or 7.7 nanomoles of Moco, blue), *Vc*MCP, (0.0077, 0.077, 0.77, 1.93, 3.85, 7.7, or 15.8 nanomoles of Moco, orange) or *Bt*XO (0.018, 0.18, 1.8, 4.38, 8.75, or 17.5 nanomoles of Moco, brown). For each experiment, animals were allowed to develop for 72 hours and animal lengths were determined. Mean and standard deviation are displayed for each data point. Sample size (n) was 10 individuals assayed for each data point. (C,D) Representative images of *moc-1(ok366) C. elegans* cultured for 72 hours on Δ*moaA E. coli* supplemented with (C) 7.7 nanomoles of Moco bound to *Vc*MCP or (D) equivalent amounts of apo-*Vc*MCP (-Moco). Blue arrowhead indicates a fertile adult, while yellow arrowheads denote animals arrested early in larval development. Scale bar is 250μm.

To test if the ability of *C. elegans* to harvest Moco from protein was more general to other Moco-binding proteins, we recombinantly expressed and purified two additional Moco-binding proteins in *E. coli*; nitrate reductase from the red bread mold *Neurospora crassa* (*Nc*NR) and Moco-carrier protein from the green algae *Volvox carteri* (*Vc*MCP) (Witte et al. 1998; Fischer et al. 2005; Hercher et al. 2020b) We also utilized the commercially available Moco-using enzyme xanthine oxidase (*Bt*XO) purified from bovine milk (Enroth et al. 2000). Each Moco-binding protein was supplemented to *moc-1* mutant *C. elegans* fed Δ*moaA* mutant *E. coli*. Similar to *Ec*YiiM, supplementation with Moco bound to either *Vc*MCP or *Nc*NR supported the growth of *moc-1* mutant *C. elegans* in the absence of any other dietary Moco (**Fig. 2)**. To a lesser extent, *Bt*XO supplementation also supported the growth of *moc-1* mutant animals cultured on Moco-deficient *E. coli* (**Fig. 2A,B)**. A possible explanation for the reduced efficacy of supplemental *Bt*XO compared to *Ec*YiiM, *Vc*MCP, or *Nc*NR might be the form of Moco that is bound by these proteins. *Bt*XO binds the xanthine oxidase form of Moco while *Ec*YiiM, *Vc*MCP, and *Nc*NR bind the sulfite oxidase form of Moco (**Fig. 1A**) (Namgung et al. 2018; Fischer et al. 2005; Hille et al. 2014; Hercher et al. 2020b). In *C. elegans* and other animals sulfite oxidase (SUOX-1) is the key Moco-requiring enzyme necessary for viability; *suox-1* null mutant animals arrest development similar to Moco-deficient animals (Warnhoff and Ruvkun 2019). We speculate that supplementation with the sulfite oxidase form of Moco can supply the appropriate Moco to support *C. elegans* SUOX-1 activity whereas the supplementation with the xanthine oxidase form of Moco cannot. Alternatively, supplementation with the sulfite oxidase form of Moco may result in the partial conversion, via Moco sulfurase (encoded by *C. elegans mocs-1*), of that supplemental Moco into the xanthine oxidase form (**Fig. 1A**). Thus, by providing the sulfite oxidase form of Moco we may be providing both forms of eukaryotic Moco making it a more effective treatment for complete Moco deficiency in *C. elegans*. Supplementation with the xanthine oxidase form of Moco would likely not result in synthesis of the sulfite oxidase form of Moco as there is no known enzyme that desulfurates the xanthine oxidase form of Moco.

To further demonstrate that the growth of *C. elegans moc-1* mutant animals was conferred by supplementation of the Moco prosthetic group and not by the supplemental purified proteins, we purified apo-*Vc*MCP from bacteria unable to synthesize Moco. Supplemental apo-*Vc*MCP did not support the growth of *moc-1* mutant *C. elegans* fed Δ*moaA* mutant *E. coli* (**Fig. 2A,C,D)**. Taken together these data demonstrate that the animal *C. elegans* is able to acquire and harvest the Moco prosthetic group when it is provided as a dietary supplement in complex with Moco-binding proteins. These proteins have diverse structures and functions and originate from both prokaryotes and eukaryotes. As such, the acquisition of protein-bound Moco by *C. elegans* is not specific to certain Moco-binding proteins and may reflect a general strategy for acquisition of functional Moco from the animals’ diet or microbiome. The remaining experiments were all performed with supplemental Moco bound to *Vc*MCP due to its well-characterized role in Moco binding and our established protocols for its production (Witte et al. 1998; Hercher et al. 2020b).

One model for the rescue of *C. elegans* Moco deficiency is that supplemental protein-bound Moco is directly ingested by *C. elegans.* Alternatively, the protein-bound Moco may first be taken up by *E. coli* which may process the Moco to then be ingested by *C. elegans.* To distinguish between these models, we grew Δ*moaA* mutant *E. coli* in lysogeny broth (LB) supplemented with Moco bound to *Vc*MCP. This Δ*moaA E. coli* was then separated from the culture medium by centrifugation, washed extensively, and fed to *moc-1* mutant *C. elegans* (“Diet B”, **Fig. 3)**. Although cultured with Moco bound to *Vc*MCP, the washed Δ*moaA E. coli* in Diet B did not support growth of *moc-1* mutant animals. Importantly, the supernatant medium from the same culture supported the growth of *moc-1* mutant *C. elegans* grown on a lawn of Δ *moaA E. coli* grown separately in LB alone (“Diet A”, **Fig. 3B)**. While we cannot completely exclude an active role for bacteria in the transfer of Moco to *C. elegans* under natural conditions, these data suggest that supplemental protein-bound Moco does not pass through a bacterial intermediate before being acquired by *C. elegans* (**Fig. 3B)**.

**Figure 3:**
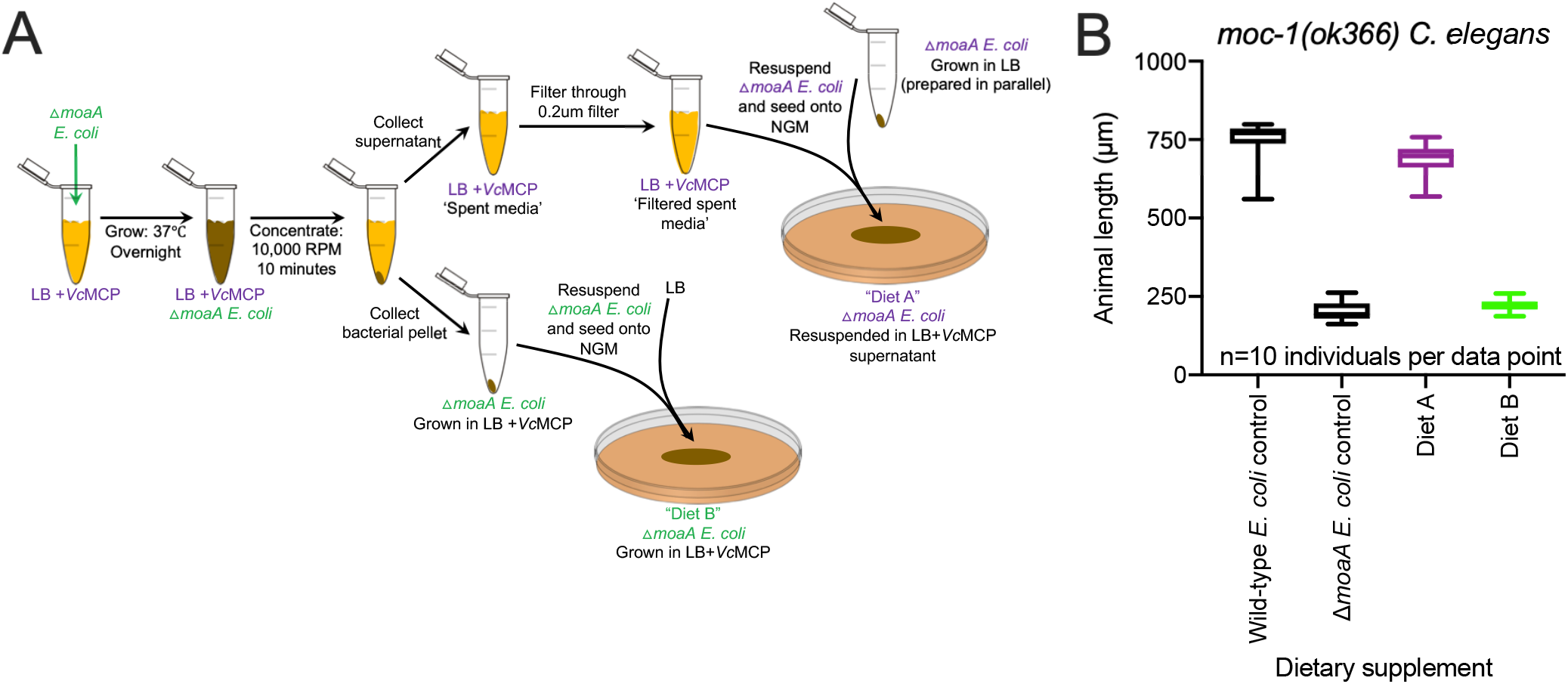
Protein-bound Moco is directly ingested by *C. elegans* (A) Experimental protocol used to generate “Diet A” and “Diet B” in Figure 3B. Δ*moaA* mutant *E. coli* were cultured overnight at 37°C in 500μl of LB supplemented with 39 nanomoles of *Vc*MCP-bound Moco. Bacterial cells were then concentrated, and the supernatant was removed for use in “Diet A”. Bacterial cells were washed, resuspended in LB, and seeded onto NGM to be fed to *moc-1(ok366) C. elegans* (“Diet B”). The supernatant from this culture (spent LB+*Vc*MCP media) was filtered (0.20μm filter, Corning) to remove remaining bacterial cells and used to resuspend a separate concentrated culture of Δ*moaA* mutant *E. coli* that was grown only in LB. This was then seeded onto NGM to be fed to *moc-1(ok366) C. elegans* (“Diet A”). (B) *moc-1(ok366)* mutant *C. elegans* were synchronized at the L1 stage and cultured on Wild-type *E. coli*, Δ*moaA E. coli,* “Diet A”, or “Diet B” (see Figure 3A for Diet A and B descriptions). For each experiment, animals were allowed to develop for 48 hours. Box plots display the median, upper, and lower quartiles while whiskers indicate minimum and maximum data points. Sample size (n) was 10 individuals assayed for each experiment.

### Moco bound to protein is stable

The instability and oxygen sensitivity of Moco has limited cell biological studies of Moco transport and precluded it from therapeutic consideration (Schwarz 2016). The *Vc*MCP-bound Moco used in “Diet A” (**Fig. 3)**was incubated at 37°C overnight, and still retained its activity and bioavailability, suggesting remarkable stability. To biochemically demonstrate the stability of Moco bound to protein, we measured the ability of mature Moco to stay in complex with *Vc*MCP, *Ec*YiiM, *Nc*NR, and *Bt*XO over time (**Fig. 4**). Free Moco is highly unstable, however it can be oxidized to ‘Form A’, a stable and fluorescent Moco-derivative that is quantifiable via HPLC (Johnson et al. 1980; Hercher et al. 2020a). Using measurements of Form A and protein concentration, we first determined the initial Moco occupancy of purified *Vc*MCP (22%) as well as *Ec*YiiM (4%), *Nc*NR (50%), and *BtXO* (50%) (**Fig. 4A,B**). We then assessed the stability of each purified Moco:protein complex by determining Moco retention over time at different temperatures (**Fig. 4C-F)**. All 4 Moco:protein complexes were remarkably stable, showing no significant protein degradation and retaining between 43 and 83% of their original Moco content after 96 hours of incubation at ambient temperature (**Fig. 4C-F**). This stability is surprising and suggests purification of protein-bound Moco as a new strategy for the production and delivery of therapeutically active Moco to treat MoCD.

**Figure 4:**
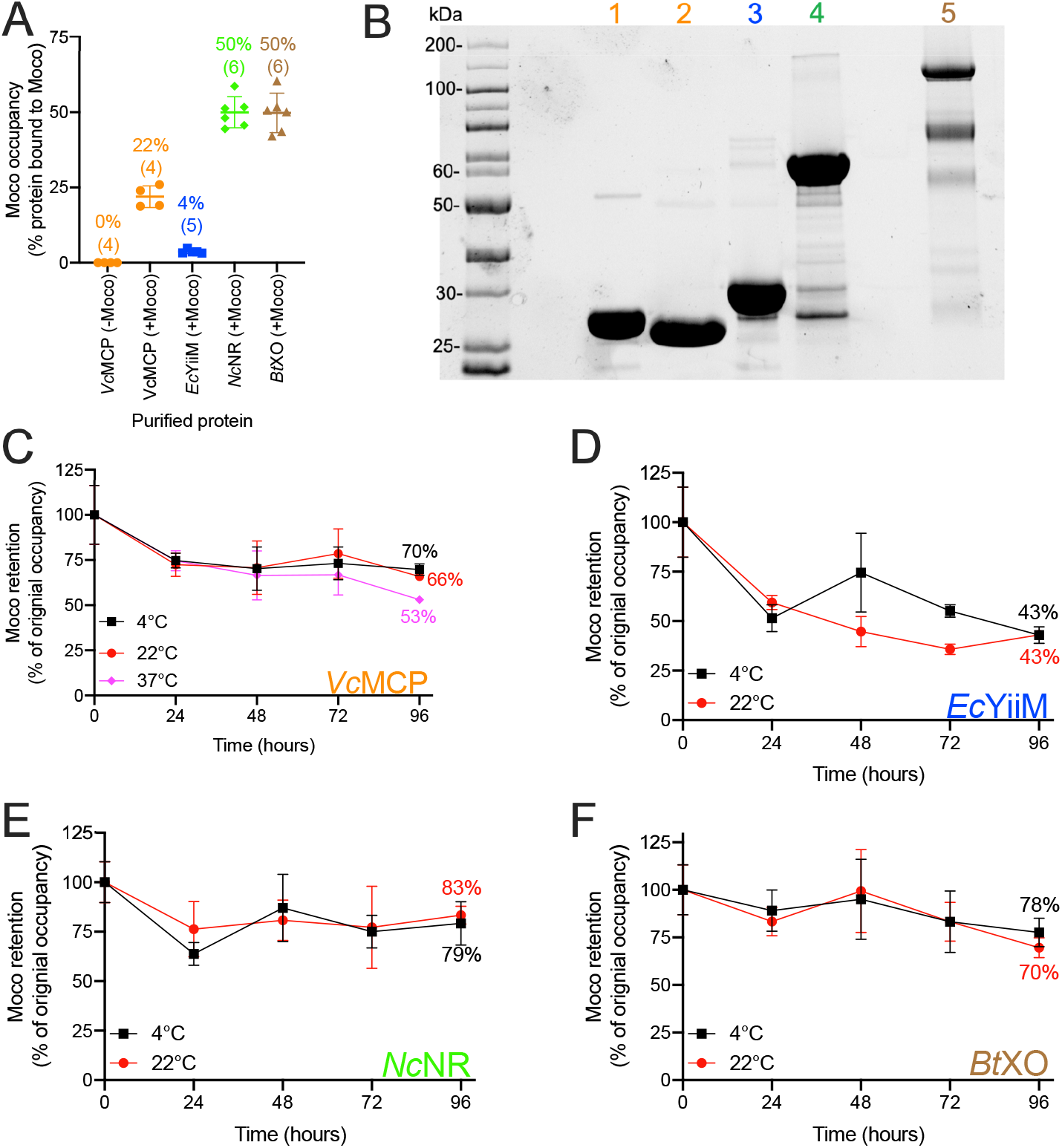
Stability of protein-bound Moco. (A) Moco occupancies for purified *Ec*YiiM, *Vc*MCP, *Nc*NR, and *Bt*XO were determined by measurements of the Moco-derivative Form A and protein concentration. Moco occupancy is the percentage of protein molecules that are bound by a Moco prosthetic group. Moco occupancy was determined for *Vc*MCP purified from Moco-producing (+Moco) and Moco-deficient (-Moco) *E. coli.* The sample size (n) is displayed for each protein and each data point is individually presented with the mean and standard deviation. (B) Protein gel demonstrating the purity of [1] *Vc*MCP purified from Moco-deficient *E. coli*, [2] *Vc*MCP, [3] *Ec*YiiM, and [4] *Nc*NR purified from Moco-producing *E. coli*, and [5] *Bt*XO purified from bovine milk. The gel displays all protein using the TGX Stain-Free system (Bio-Rad). (C-F) The amount of stable Moco retained by (C) *Vc*MCP, (D) *Ec*YiiM, (E) *Nc*NR, and (F) *Bt*XO was determined over 96 hours at 4°C (black) or 22°C (red). Moco retention of *Vc*MCP was also assessed at 37°C (pink). The Y-axis displays the Moco retention as a percentage of the original Moco occupancies (time 0) presented in Figure 4A. The sample size (n) is 3-6 replicates per protein and time point. The mean and standard deviation are displayed.

### Bioavailability of recombinant protein-bound Moco does not depend on known Moco-biosynthetic enzymes in *E. coli* or *C. elegans*

We tested if the Moco-biosynthetic enzymes are necessary for the harvesting or transport of supplemental protein-bound Moco using mutants in the dietary *E. coli*. We tested *moc-1* mutant *C. elegans* growth on wild-type bacteria, or mutant bacteria lacking the genes necessary for Moco biosynthesis. *moc-1* mutant animals were grown on mutant *E. coli* with and without supplemental Moco bound to *Vc*MCP. *moc-1* mutant *C. elegans* grew well on wild-type *E. coli* but displayed larval arrest on all 10 *E. coli* mutants defective in Moco biosynthesis (**Supplemental Fig. S1)**. Supplemental Moco bound to *Vc*MCP supported growth and development of *moc-1* mutant *C. elegans* on all 10 of the Moco-biosynthetic mutant *E. coli* demonstrating that none of these *E. coli* genes were necessary for bioavailability of supplemental protein-bound Moco (**Supplemental Fig. S1)**.

Alternatively, we speculated that the Moco-biosynthetic machinery of *C. elegans* might play a role in the bioavailability of supplemental protein-bound Moco. To test this, we used established *C. elegans* mutants in the Moco-biosynthetic pathway (**Fig. 1A**). Each of these *C. elegans moc* mutants was cultured on wild-type *E. coli*, Δ*moaA E. coli,* or Δ*moaA E. coli* supplemented with Moco bound to *Vc*MCP. All of the *moc* mutant animals grew well on wild-type bacteria and arrested growth on Δ*moaA E. coli* (**Supplemental Fig. S2A-E**). Each *C. elegans moc-*mutant displayed dramatically improved growth on Δ*moaA E. coli* when their diet was supplemented with Moco bound to *Vc*MCP (**Supplemental Fig. S2A-E**). These results demonstrate that *moc-5, moc-4, moc-3, moc-2,* and *moc-1* are not required for the bioavailability of supplemental protein-bound Moco. Thus, the machinery that facilitates Moco transport is distinct from the canonical Moco-biosynthetic pathway.

### Supplemental protein-bound Moco supports the activity of *C. elegans* SUOX-1

The lethality associated with Moco deficiency in *C. elegans* and humans is due to inactivity of sulfite oxidase (SUOX-1), a mitochondrial Moco-requiring enzyme that oxidizes the lethal toxin sulfite to sulfate. Like Moco-biosynthesis, sulfite oxidase is essential in both *C. elegans* and humans (Mudd et al. 1967; Warnhoff and Ruvkun 2019). Thus, to rescue development of otherwise Moco-deficient *C. elegans,* supplemental protein-bound Moco must be incorporated into and support the activity of *C. elegans* SUOX-1. To demonstrate this, we utilized the hypomorphic *suox-1* allele *gk738847* (D391N) (Thompson et al. 2013). Aspartic acid 391 of sulfite oxidase is highly conserved and is present in *C. elegans* and humans. The SUOX-1 D391N amino acid substitution causes partial SUOX-1 loss-of-function that is enhanced when dietary Moco is absent. Growing *suox-1(gk738847)* mutant *C. elegans* on Moco-deficient *E. coli* causes a severe developmental delay compared to its growth on wild-type Moco-producing *E coli* (**Supplemental Fig. S2F**) (Warnhoff and Ruvkun 2019). Importantly, *suox-1(gk738847)* mutant animals are wild type for their endogenous Moco biosynthetic pathway and are able to synthesize Moco *de novo*. This result shows that *suox-1(gk738847)* mutant *C. elegans* depend on both endogenous Moco biosynthesis as well as dietary sources of Moco to fully support the activity of mutant SUOX-1 D391N protein.

We hypothesized that supplemental Moco bound by *Vc*MCP would improve the viability of *suox-1(gk738847)* animals grown on Δ*moaA E. coli.* To test this, we cultured *suox-1(gk738847)* mutant animals on wild-type *E. coli*, Δ*moaA E. coli,* and Δ*moaA E. coli* supplemented with Moco bound to *Vc*MCP. Consistent with our rescue of *C. elegans* Moco deficiency, supplemental protein-bound Moco improved the growth of *suox-1(gk738847)* animals grown on Moco-deficient *E. coli* (**Supplemental Fig. S2F)**. These results suggest exogenous protein-bound Moco is absorbed, harvested, distributed to requisite cells, and re-inserted into the *C. elegans* SUOX-1 enzyme. Uncovering the cellular mechanisms that facilitate these processes is an important goal of future research. Pathways for absorption and distribution of heme and vitamin B12 serve as useful paradigms for the possible mechanisms of Moco transport. These metal-containing prosthetic groups require chaperones (Dieckgraefe et al. 1988; Chen et al. 2011), receptors (Moestrup et al. 1998), and transporters (Rajagopal et al. 2008) to facilitate their movement among cells and tissues. We anticipate similar systems will coordinate the transport of Moco in *C. elegans* and humans.

Our data demonstrate the ability of an essential protein-packaged prosthetic group to cross cell membranes. We show that this transfer naturally occurs between multiple organisms and among the cells and tissues of a single organism. Because Moco biosynthesis and utilization is as ancient as the last universal common ancestor, the unknown Moco-transport pathway is likely to be general to all animals. Furthermore, roughly 70% of bacterial genomes encode Moco-biosynthetic enzymes making the intestinal microbiome a potential reservoir for this cofactor (Zhang and Gladyshev 2008). Similarly, the human diet might also be a source of exogenous protein-bound Moco as most plants and animals synthesize and utilize Moco. Our results with the nematode *C. elegans* may stimulate future exploration of the therapeutic potential of protein-bound Moco (from dietary, microbiome, or recombinant sources) for the treatment of MoCD.

## Materials and Methods

### General methods and strains

*C. elegans* strains were cultured at using established protocols (Brenner 1974). The wild-type strain of *C. elegans* was Bristol N2. *C. elegans* mutant strains used in this work are listed here. LGI: GR2253 *moc-4(ok2571).* LGIV: MH3266 *moc-3(ku300).* LGV: GR2255 *moc-2(mg595).* LGX: GR2254 *moc-1(ok366),* GR2256 *moc-5(mg589),* and GR2269 *suox-1(gk738847*).

*E. coli* strains were cultured using standard methods. The wild-type strain of *E. coli* was BW25113, the parental strain of the Keio *E. coli* knockout collection (Baba et al. 2006). The *E. coli* mutants used in this work were JW0764 (Δ*moaA::*Kan^r^), KJW1 (Δ*mog*), KJW2 (Δ*moaA*), KJW3 (Δ*moaC*), KJW4(Δ*moaD*), KJW5(Δ*moaE*), KJW6(Δ*moeB*), KJW7(Δ*moeA*), KJW8(Δ *modA*), KJW9(Δ*modC*), and KJW10(Δ*ydaV*) (Baba et al. 2006; Warnhoff and Ruvkun 2019). Strains KJW1-KJW10 were only used to produce the data in Supplemental Fig. S1. JW0764 was used in all other experiments withΔ*moaA E. coli*.

### *C. elegans* growth assays

*C. elegans* were synchronized at the first stage of larval development (L1) then cultured on NGM seeded with wild-type or mutant *E. coli.* For some experiments, growth conditions were supplemented with various forms and amounts of protein-bound Moco. For experiments with supplemental protein-bound Moco, we report the total amount of Moco added to the *C. elegans* growth conditions and assume equal protein diffusion through the solid agarose media during the experiments. *C. elegans* animals were allowed to grow and develop for 48 or 72 hours (specified in Figure Legends) at 20°C. Sample size (n) is individual animals measured and is reported in the Figures and Figure Legends.

Live animals were imaged using an Axio Zoom.V16 microscope (Zeiss) equipped with an ORCA-Flash4.0 digital camera (Hamamatsu). Images were captured using ZEN software (Zeiss) and processed with ImageJ (NIH). Animal length was measured from the tip of the head to the end of the tail. The median and upper and lower quartiles were calculated using GraphPad Prism software.

### Purification and characterization of Moco-binding proteins

Moco-binding proteins were expressed and purified using standard methods (Hercher et al. 2020b). The full-length *yiiM* coding sequence was amplified from *Escherichia coli* DH5α (*Ec*YiiM) and the full-length Moco carrier protein coding sequence from *Volvox carteri* (*Vc*MCP) was synthesized and codon optimized for *E. coli* (Hercher et al. 2020b). The coding sequence for *Neurospora crassa* nitrate reductase (*Nc*NR) was shortened to include only the Moco-binding and dimerization region (amino acids 113-592). The coding sequences for *Ec*YiiM, *Vc*MCP, and *Nc*NR were inserted into the pONE-CP plasmid, producing proteins fused to a C-terminal Streptavidin tag. Streptavidin-tagged proteins were expressed using the *E. coli* strain TP1000 which accumulates Moco due to a deletion in the Mob operon (Palmer et al. 1996). As a negative control, *Vc*MCP was also purified from the *E. coli* strain RK5204 which is unable to produce Moco (Stewart and MacGregor 1982). Bovine xanthine oxidase (*Bt*XO) was purchased from Sigma-Aldrich (X1875, batch SLCB1289).

Protein concentrations were determined using absorption at 280nm and the Pierce BCA Protein-Assay (Thermo Scientific). Absorption was measured using a Multiskan GO Microplate Spectrophotometer (Thermo Scientific). Quantification of Moco content of the proteins was conducted using HPLC-based measurements of Form A, a stable and fluorescent Moco-oxidation product (Hercher et al. 2020a). Stability of protein-bound Moco was assessed by incubating the Moco:protein complexes at various temperatures (4°C, 22°C, and 37°C) for 96 hours. In 24-hour intervals, protein samples were centrifuged at 4°C to remove precipitated protein and protein concentration and Moco content were determined as described above.

## Acknowledgments

We thank the *Caenorhabditis* Genetics Center (CGC) for providing *C. elegans* strains and the National BioResource Project (NIG, Japan) for providing the Keio *E. coli* knockout collection. We thank Dr. Jörn Krausze for help in crystal structure-guided determination of the stereochemistry of Moco intermediates in Figure 1A. This work was funded by an NIH Grant (5R01GM044619-26) to G.R., a DFG grant (GRK2223/1) to R.R.M., and a Damon Runyon Fellowship (DRG-2293-17) to K.W.

## Supplemental Material for

**Supplemental Figure 1:**
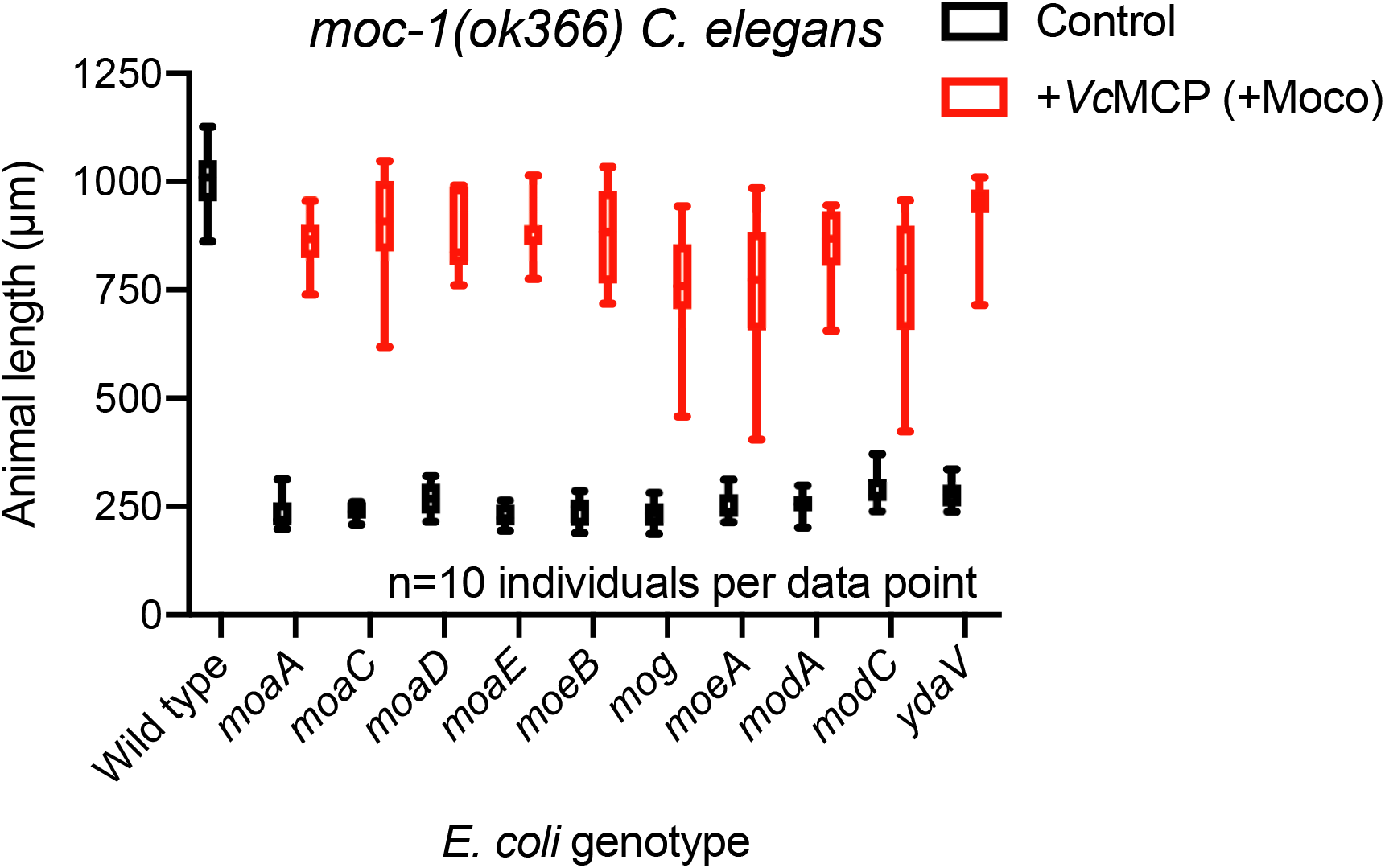
*E. coli* Moco-biosynthetic enzymes are not necessary for uptake of supplemental protein-bound Moco by *C. elegans*. *moc-1(ok366)* mutant *C. elegans* were synchronized at the L1 stage and cultured on wild-type or mutant *E. coli* for 72 hours. Each growth assay performed with mutant *E. coli* was done with or without 3.3 nanomoles of supplemental Moco bound to *Vc*MCP. The Y axis shows animal length (μm), where 1,000μm roughly corresponds with fertile adulthood and 250μm roughly corresponds to the L1 stage. Box plots display the median, upper, and lower quartiles while whiskers indicate minimum and maximum data points. Sample size (n) was 10 individuals assayed for each experiment.

**Supplemental Figure 2:**
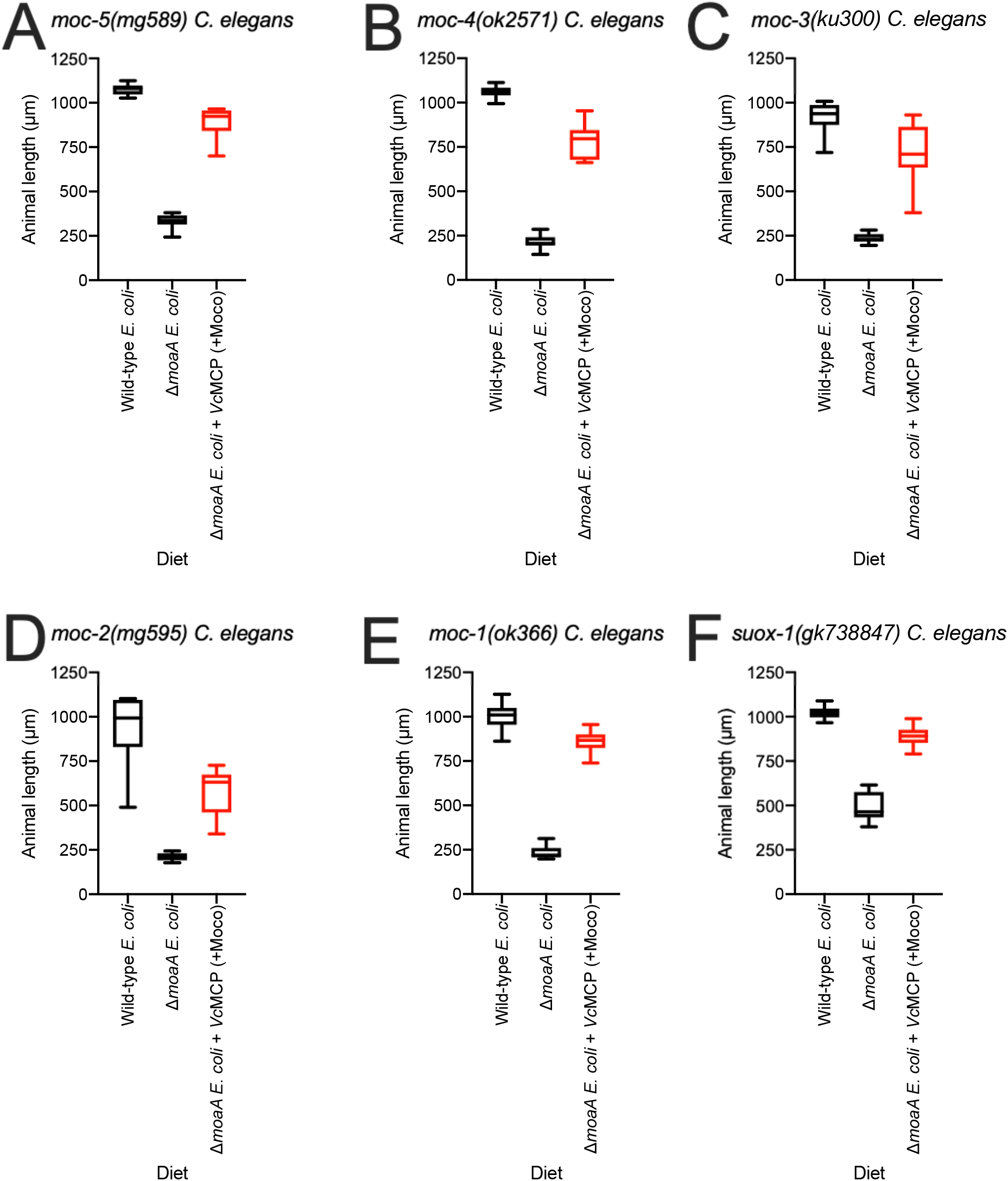
*C. elegans* Moco-biosynthetic enzymes are not necessary for uptake of supplemental protein-bound Moco. (A) *moc-5(mg589),* (B) *moc-4(ok2571),* (C) *moc-3(ku300),* (D) *moc-2(mg595),* (E) *moc-1(ok366)* and (F) *suox-1(gk738847)* mutant animals were cultured from synchronized L1 larvae for 72 hours on wild-type, Δ*moaA E. coli,* or Δ*moaA E. coli* supplemented with 3.3 nanomoles of Moco bound to *Vc*MCP. The Y axis shows animal length (μm), where 1,000μm roughly corresponds with fertile adulthood and 250μm roughly corresponds to the L1 stage. Box plots display the median, upper, and lower quartiles while whiskers indicate minimum and maximum data points. Sample size (n) was 10 individuals assayed for each experiment.

